# Potential of imputation for cost-efficient genomic selection for resistance to *Flavobacterium columnare* in rainbow trout (*Oncorhynchus mykiss*)

**DOI:** 10.1101/2023.01.04.522693

**Authors:** C. Fraslin, D. Robledo, A. Kause, R.D. Houston

## Abstract

**Background:** *Flavobacterium columnare* is the pathogen agent of columnaris disease, a major emerging disease affecting rainbow trout aquaculture. Selective breeding using genomic selection has potential to achieve cumulative improvement of host resistance. However, genomic selection is expensive partly due to the cost of genotyping high numbers of animals using high-density SNP arrays. The objective of this study was to assess the efficiency of genomic selection for resistance to *F. columnare* using *in silico* low-density (LD) panels combined with imputation. After a natural outbreak of columnaris disease, 2,874 challenged fish and 469 fish from the parental generation (n=81 parents) were genotyped with 27,907 SNPs. The efficiency of genomic prediction using LD-panels was assessed for panels of 10 different densities, created *in silico* using two sampling methods, random and equally spaced. All LD-panels were also imputed to the full 28K HD-panel using the parental generation as the reference population, and genomic predictions were reevaluated. The potential of prioritizing SNPs showing association with resistance to *F. columnare* was also tested for the six lower densities.

**Results:** Similar results were obtained with random and equally spaced sampling of SNPs for accuracy of both imputation and genomic predictions. Using LD-panels of at least 3,000 makers or lower density panels (as low as 300 markers) combined with imputation resulted in comparable accuracy to the 28K HD-panel and 11% higher accuracy than pedigree-based predictions.

**Conclusions:** Compared to using the commercial HD-panel, LD-panels with imputation may provide a more affordable route to genomic prediction of breeding values, supporting wider adoption of genomic selection in aquaculture breeding programmes.

## INTRODUCTION

Aquaculture production has increased dramatically over the past decades and is now supplying more aquatic products than fisheries (1). Compared to livestock production, the domestication of most aquaculture species is recent and not all species benefit from a modern selective breeding programme (2). Nonetheless, selective breeding has been successfully implemented for an important number of aquaculture species, and the recent development of high throughput genotyping technologies, such as single nucleotide polymorphism (SNP) arrays, has opened the gate for the implementation of genomic selection for the most important species (2–4). Genomic selection uses genome-wide marker information (mainly SNPs), used to generate genomic relationship matrices, to predict the breeding value of genotyped selection candidates based on genotype and phenotype information gathered in a reference population (5,6). In aquaculture breeding programmes, many traits under selection cannot be measured directly on the candidates (e.g. fillet yield or disease resistance traits), and thus are measured on their full and half-sibs (7). These so-called sib traits are the perfect target for the implementation of genomic selection because it captures the within-family genetic variation in addition to the between-family genetic variation. Over the recent years, a large number of studies have demonstrated that the application of genomic selection significantly improves the response to selection of aquaculture breeding programmes (2,8–11).

The late implementation of genomic selection in aquaculture breeding programmes compared to terrestrial livestock species was partly due to the lack of high throughput genotyping platforms for most species and due to the significant cost of genotyping the large number of individuals required for efficient genomic selection. Therefore, so far, genomic selection has only been implemented for a handful of aquaculture species with the largest production value, and typically by the largest companies. Several strategies have been investigated to reduce the genotyping cost and make genomic selection more affordable for small and medium scale breeding programmes, such as genotyping only a proportion of the individuals (12,13), DNA pooling to build a reference population (14–16) or the use of medium and low-density SNP panels that are typically cheaper to produce.

To date, many studies have investigated the potential use of low-density (LD) SNP panels for genomic selection in various aquaculture species and they concluded that LD-panels containing 1,000-2,000 SNPs (17,18) to 6,000 SNP (19). Depending on the species and the trait (reviewed in Song et al., 2022) (11) are sufficient to achieve an accuracy of genomic prediction similar to that obtained with a medium or high-density panel. In those studies, further reduction of the density to hundreds of SNPs resulted in a significant drop in accuracy (17,20). This issue could potentially be resolved via the use of imputation. Imputation predicts the missing genotypes in a LD-genotyped target population using information from a HD-genotyped reference population. Imputation relies on linkage disequilibrium information in a population-based imputation approach, or on linkage information in a family-based imputation approach (21). The usefulness of imputation in genomic prediction has been studied for various farmed crops and animals (22–25) and is now implemented on a routine basis in cattle genomic selection. A few recent studies have investigated the impact of imputing low density to medium or high density genotypes on the accuracy of genomic prediction in major aquaculture species such as Atlantic salmon, rainbow trout and tilapia (26–33). Those studies showed that a cost-efficient genomic selection could be achieved with a combined approach of low-density genotyping and imputation.

One example of a programme where the benefits of such cost-efficient genotyping approaches may be realised is the Finnish rainbow trout breeding programme. It was established in the late 1980s and it relies upon pedigree-based information obtained by initial rearing of families in separate tanks until the fish are big enough to be tagged and pooled together in larger tanks (34,35). In recent years, columnaris disease (CD) caused by *Flavobacterium columnare*, has become a major concern for rainbow trout farming in Finland. *Flavobacterium columnare* is a bacterium distributed worldwide and affecting fresh water fish in warm water temperatures, usually above 18-20°C, but has also been reported to affect salmonids in cooler water (36–39). *F. columnare* causes acute and chronic infections which mainly manifest as tissue and gill necrosis especially in small fish, leading to high mortality if the disease is not treated (37,39,40). In a recent study on resistance to *F. columnare* in two Finnish rainbow trout populations, we detected genetic variation and quantitative trait loci (QTL) associated with resistance to this disease and advocated for the use of genomic selection (and / or marker assisted selection) to improve this trait (41,42). Implementing genomic selection may speed up genetic gain for various traits including resistance to CD, but to date the cost of genotyping remains prohibitive for the Finnish breeding programme. The aim of this study was to assess the efficiency of genomic selection to improve rainbow trout resistance to *F. columnare* using low-density SNP panels created *in silico* and imputation using three different SNP selection strategies: i) randomly sampled along chromosomes, ii) equally spaced within each chromosome and iii) most significant SNPs based on genome-wide association study (GWAS) results.

## Materials & Methods

### Ethical statement

The establishment of progeny families at Luke’s research facilities followed the protocols approved by the Luke’s Animal Care Committee, Helsinki, Finland. Hanka-Taimen Oy, a fish farming company, has authorisation for fish rearing and experiments, and both parties comply with the EU Directive 2010/63/EU for animal experiments.

### Fish rearing and disease outbreak management and genotyping

The fish used in this study come from the Finnish national breeding programme for rainbow trout, managed by the Natural Resources Institute Finland (LUKE). Fish rearing, phenotyping and genotyping have been described by Fraslin et al., (2022) (41). Briefly, in May 2019, 81 rainbow trout breeding candidates [33 females (dams) and 48 males (sires)] were selected among 567 fish from the Finnish national breeding programme, based on their relationships and genetic contribution to maintain a predetermined inbreeding coefficient of less than 1% per generation. The 33 dams and 48 sires were mated to create 105 full-sib families. A 0.5 dL of eggs from each mating were pooled after fertilisation and incubated together.

In June 2019, around 30,000 fry were separated into three fingerling tanks, resulting about 100 fish per family per tank, at a multiplier farm of Hanka-Taimen Oy (Finland) that uses water from a nearby stream with naturally occurring CD outbreaks. From arrival at this multiplier farm (considered as day 0 of the study), the fish mortality and any disease signs were monitored twice a day. On day 11 of the experiment, fish in all three tanks started to show signs of CD (saddleback lesions), seven dead or dying fish were sampled and sent to a veterinarian to confirm the CD diagnosis. The presence of the pathogen was confirmed by PCR. From day 20 to 24, a piece of tail was taken, for later DNA extraction, from 510 fish per tank, randomly chosen amongst the dead or dying fish with clear CD signs (considered as susceptible). At day 26, the three tanks were treated following the veterinarian guidelines against *F. columnare* with an approved treatment of salt, chloramine and medical feed until day 32. On the last day of the experiment, day 99, a piece of tail was collected, for latter DNA extraction, on about 506 fish per tank, randomly sampled among the fish still alive at that time (considered as resistant). A total of 3,057 challenged fish (1,538 susceptible and 1,519 resistant) and 570 fish from the parental generation (including the 81 parents) were genotyped using the 57K SNP Axiom^™^ Trout Genotyping Array (43). The genotypes of all 3,624 individuals were called together in a single run using Axiom Analysis Suite software (v.4.0.3.3) with the recommended standard SNPs quality controls. Only SNP classified as “highly polymorphic” by the software were kept for further quality control (n=36,020 SNPs). The software plink (v.1.9) (44) was used to perform quality control on SNPs and individuals based on deviation from Hardy-Weinberg equilibrium (*p-value* ≤ 10^-6^), minor allele frequency (≥ 0.05), marker call rate (≥ 0.95) and individual call rate (≥ 0.9). The final dataset comprised 2,874 challenged fish and 469 fish from the parental generation (including 78 parents of the challenged fish), all genotyped for 27,907 SNPs. Those 28K SNPs were considered as the high-density (HD) panel for the remaining of the analysis.

### In silico low-density panels

The impact of reducing the SNP density on genomic prediction was tested with low-density (LD) SNP panels created *in silico* using three different sampling methods. For the first method, SNPs to be included in the LD-panels were sampled randomly within each chromosome, with the number of SNPs sampled from a given chromosome being proportional to the physical length of the chromosome in the *O. mykiss* reference genome (Omyk_0.1) (45). This random selection method will be referred to as RandLD (Random Low-Density). For the second sampling method, referred to as EquaLD (Equally spaced Low-Density), SNPs were selected so that they would be equally spaced within each chromosome. For those 2 methods we used the CVrepGPAcalc package (26) to create 10 different panels for each of the 10 densities (300; 500; 700; 1,000; 3,000; 5,000; 7,000; 10,000; 15,000 and 20,000 SNPs). Replicates were allowed to overlap by chance and the final number of SNPs within each panel was allowed to slightly vary from the target density. On average, the RandLD-panels created contained between 10 and 15 SNPs more than the target density (Supplementary Table S1). All the replicates for each density contained the same number of SNPs. The EquaLD-panels were more variable and contained, on average, between 1 and 2 SNPs less than the target density for LD panels from 300 to 1K and, on average, between 2 and 104 SNPs more for densities from 3K to 15K. For both methods, the 20K LD-panel contained about 1K less SNPs than targeted with 19,803 SNPs for the RandLD panels and on average 19,039 SNPs (± 55 SNPs) for the EquaLD-panels (Supplementary Table S1).

Finally, the outcome of GWAS for resistance to *F. columnare* was used in another SNP sampling method to create Top Low-Density (TopLD) panels based on the *p*-value estimated by the GWAS in order to see the effect of including SNPs with a significant effect on resistance into the LD-panel. The SNP effect and *p*-value was computed in a GWAS performed with a mixed linear model association (mlma) with the leave-one-chromosome-out (loco) option implemented in GCTA (46) using the model presented in Fraslin et al., (2022) (41) with resistance analysed as a binary trait (0=alive, 1=dead) and the tank number included as a fixed effect. One of the pitfalls of the creation of those TopLD panels is that estimating the SNP effects in a GWAS that includes the whole population and then validating those SNP effects on a sub-sampling of the population in a genomic prediction approach (validation group) can lead to inflated prediction accuracies. In order to avoid this issue and to have the SNP effects estimated in an independent group from the validation set, we used the same “leave-one-group” out approach in a five folds cross validation scheme as defined for the evaluation of genomic prediction accuracy (see below) to perform 100 independent GWAS. Specifically, the fish were randomly separated into 5 groups, one of them (20% of the population, validation set) was excluded from the analysis and the phenotype and genotype information of the remaining 80% of the fish (training set) was used in the GWAS to estimate each SNP effect and *p*-value for association to resistance, and this was repeated for the 5 groups. This was replicated 20 times to match the 20 replicates of the genomic prediction evaluation, so a total of 100 GWAS were performed. The SNPs were then ranked, within each GWAS, from lowest *p*-value (most significant association) to highest *p*-value (least significant association) and the N first SNPs were sampled to create a TopLD-panel. A total of 100 TopLD-panels were created for each density and we tested 6 different densities composed of the best 300, 500, 700, 1000, 3000 and 5000 SNPs. Since the SNPs were selected based on the GWAS results not all chromosomes were represented in the lower density TopLD-panels, and chromosomes 3 and 5 were over-represented due to the presence of major QTL associated with resistance to *F. columnare* (41,42).

### Imputation of low-density panels to a high density of 27,970 SNPs

Imputation was performed only for the RandLD and EquaLD panels using FImpute3 software (47). LD genotypes from the offspring were imputed back to the full ^~^28K SNPs using a combined population and pedigree-based imputation method with the HD-genotyped parents (n=469 fish) as the reference population. The “parentage_test” option was used with an error rate threshold of 0.05 (“/ert mm 0.05”) to find progeny-parent mismatches based on the pedigree, and in case of Mendelian inconsistency between progeny and parents for non-missing genotypes the original genotypes were kept intact using the option “keep_og”. The accuracy of imputation was estimated as the Pearson correlation coefficient between true and imputed genotypes for only the SNPs that were removed to create the LD-panels. After imputation, another quality control was performed and imputed makers with a maf below 0.05 were removed.

### Genomic evaluation of low-density SNP panels before and after imputation

The (genomic) estimated breeding values [(G)EBV] of fish were computed using mixed linear BLUP animal model based on pedigree (PBLUP) or genomic (GBLUP) information using the BLUPF90 software (48). Disease resistance was analysed as a binary trait (0 = alive; 1 = dead) with the rearing tank as a fixed effect in the statistical model. The efficiency of genomic prediction was estimated by a 5-fold cross validation procedure using Monte-Carlo “leave-one-group-out” method. The phenotypes of 20% of the fish (validation set) were masked, and their (G)EBVs were predicted using the phenotype and genotype information of the remaining 80% fish (training set). This procedure was repeated 20 times for the PBLUP, and the GBLUP with all 28K SNPs (HD-GBLUP) and for each of the 10 replicates of both the RandLD and EquaLD panels, pre- and post-imputation. For the TopLD panels, the genomic prediction was only performed for the un-imputed panels, and as the groups created for the cross-validation procedure were the same as the ones to select the SNPs in the panels, the performance of each TopLD panel was only tested within its corresponding validation set.

The performance of genomic prediction was assessed by estimating the accuracy of genomic prediction and the estimation bias (49). The accuracy was computed, for each SNP panel, as the mean over the 100 replicates of the correlation between the (G)EBV and the true phenotype of the fish in the validation group, divided by the square root of the genomic-based heritability (h^2^ = 0.21 as estimated in Fraslin et al., 2022) (41). The estimation bias was computed, for each SNP panel, as the regression coefficient of the true phenotype (on y-axis) on the (G)EBVs (on x-axis). This coefficient is a measure of the degree of inflation and is expected to be equal to 1 in the absence of bias, a value below 1 represents an over-dispersion of (G)EBVs and a value above 1 represents an under-dispersion of (G)EBVs (50).

## Results

### Genomic prediction with LD-panels

Accuracy of the PBLUP and HD-GBLUP were previously estimated in (41). The pedigree-based prediction accuracy was estimated as 0.59 (± 0.080 sd) and the GBLUP genomic evaluation using the HD panel increased the prediction accuracy by 14% (0.68 ± 0.076).

Decreasing the number of SNPs decreased the accuracy of genomic prediction (Figure 1), and no significant difference was observed between the random or equally spaced methods of SNP sampling. For both RandLD and EquaLD, prediction accuracies obtained with 300 to 500 SNPs were close to the accuracy obtained with the pedigree-based analysis. Encouragingly, prediction accuracies obtained with LD-panels from 7,000 SNPs and above were close to the accuracy obtained with the HD-panel (−1% in accuracy compared with the HD-GBLUP). Accuracies obtained with 1,000 SNPs were only 4% higher than those obtained with the pedigree, whereas the accuracy obtained with only 3,000 SNPs was 3% lower than the accuracies obtained with the HD-panel and thus 11% higher than the accuracy obtained with the pedigree only.

**Figure 1.**
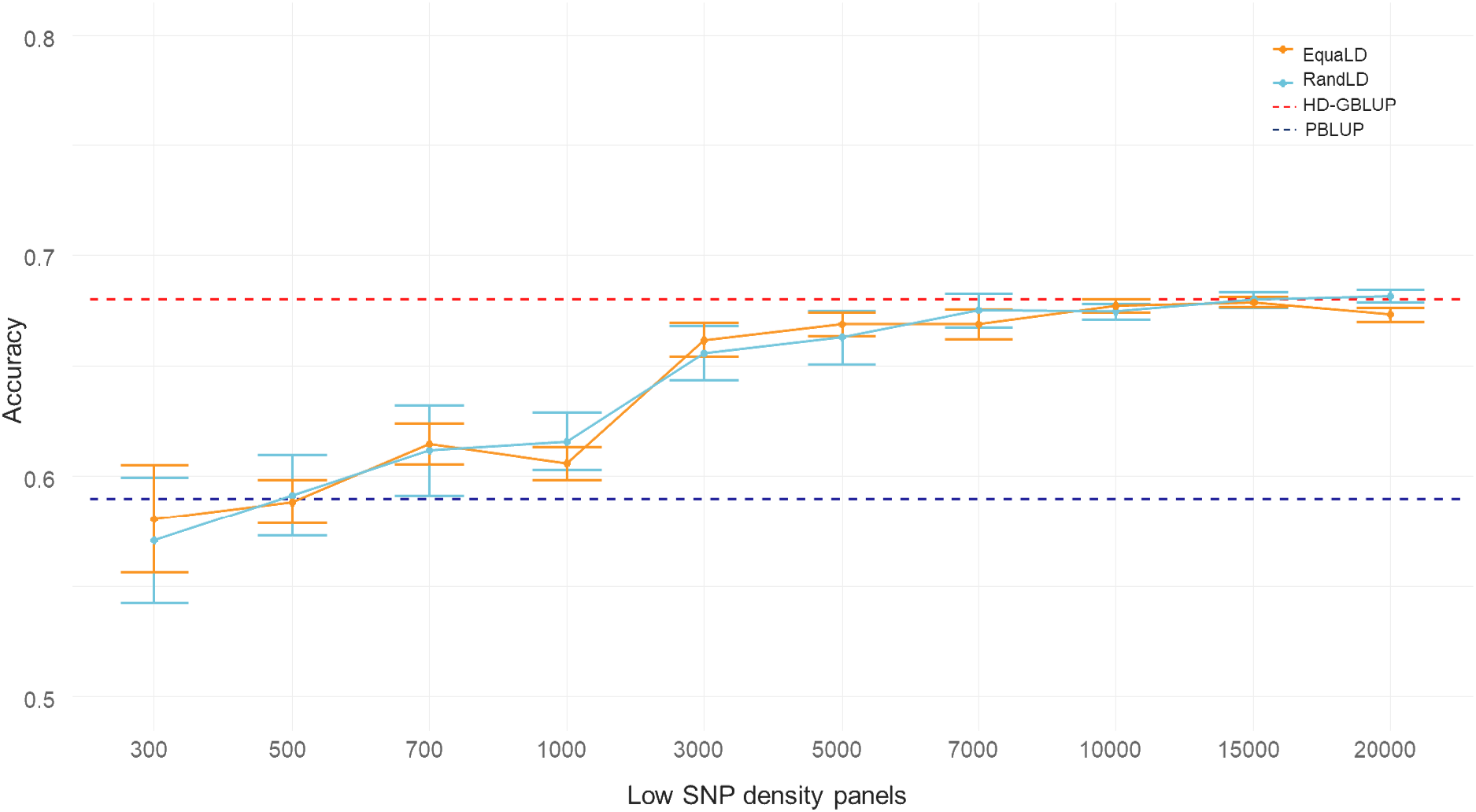
Accuracy of genomic prediction for resistance to *F. columnare* in rainbow trout, obtained with different low-density SNP panels (no imputation). The horizontal red dotted line is the average accuracy for the HD-GBLUP (28K) prediction (0.68), the horizontal blue dotted line is the average accuracy for the pedigree-based BLUP prediction (0.59). The light blue line is the accuracy obtained with the Random SNP sampling LD-panels (RandLD). The orange line is the accuracy obtained with the equally spaced SNP sampling LD-panels (EquaLD). The mean (dots) and standard deviations (bars) are taken from 10 replicates of each marker density.

Variation among the 10 replicates was higher for lower densities with a bigger standard deviation (Figure 1). With 300 SNPs, the highest accuracy obtained with one panel was 0.63 for RandDL and 0.61 for EquaLD, which was similar to the average accuracy obtained with 700 or 1,000 SNPs. The lowest accuracy obtained with 300 SNPs was 0.53 for RandLD-panel and 0.55 for EquaLD-panel which is significantly lower than the accuracy obtained with the pedigree-based analysis.

The accuracy and bias of genomic prediction obtained with the TopLD panels are presented in Table 1. For the lowest densities (300 – 1,000) the accuracy of prediction obtained with SNPs selected based on their GWAS *p*-value (TopLD) was significantly better than the accuracy obtained for the same density when SNPs were selected randomly or equally spaced, except for EquaLD vs TopLD at 700 SNP were the difference was non-significant (*p*-value=0.059, Wilcox test). For higher densities (3K and 5K), prioritising the SNPs based on the GWAS significantly decreased the accuracy of genomic prediction compared to RandLD or EquaLD panels of the same densities (*p*-value ranging from 0.002 to 0.05 for EquaLD 5K and RandLD 3K, respectively).

**Table 1.** Performance of low-density SNP panels with the most significantly associated SNPs for resistance to *F. columnare* (TopLD strategy).

| SNP density | Accuracy (mean ± sd) | Bias (mean ± sd) |
| --- | --- | --- |
| **300** | 0.62 ± 0.072 | 0.59 ± 0.078 |
| **500** | 0.66 ± 0.161 | 0.60 ± 0.143 |
| **700** | 0.63 ± 0.074 | 0.57 ± 0.079 |
| **1000** | 0.63 ± 0.074 | 0.56 ± 0.078 |
| **3000** | 0.64 ± 0.072 | 0.56 ± 0.074 |
| **5000** | 0.65 ± 0.074 | 0.57 ± 0.077 |

Values are the mean accuracy and mean bias obtained as average of the 100 replicates (5 groups \* 20 replicates).

For all the TopLD panels, the GEBVs obtained were highly biased with on average a bias of 0.575, which represents an over-dispersion of the breeding values. In contracts, the RandLD and EquaLD panels showed very little bias (Supplementary Table S2).

### Imputation from low-density genotypes to 28K SNPs

The imputation accuracies for both SNP selection methods are presented in Figure 2. There was no significant difference in the accuracy of imputation for the two SNP selection methods (random vs. equidistant), expect for the 20K SNP panels where the EquaLD panel had a lower imputation accuracy. Accuracy of imputation increased rapidly from 0.58 (± 0.004) for the 300 SNPs EquaLD-panel to 0.68 (± 0.005) for the 500 SNPS EquaLD-panel, and reached a plateau at around 0.86 - 0.89 from the 7000 SNPs LD-panels. The last important increase in imputation accuracy occurred between 1,000 (0.77 ± 0.003) and 3,000 SNPs (0.83 ± 0.003), and after this density imputation accuracy increased at a lower and more steady rate. There was a drop in the imputation accuracy at 20K SNPs for the EquaLD-panel only but with a higher variability among panels (0.87 ± 0.014 sd).

**Figure 2.**
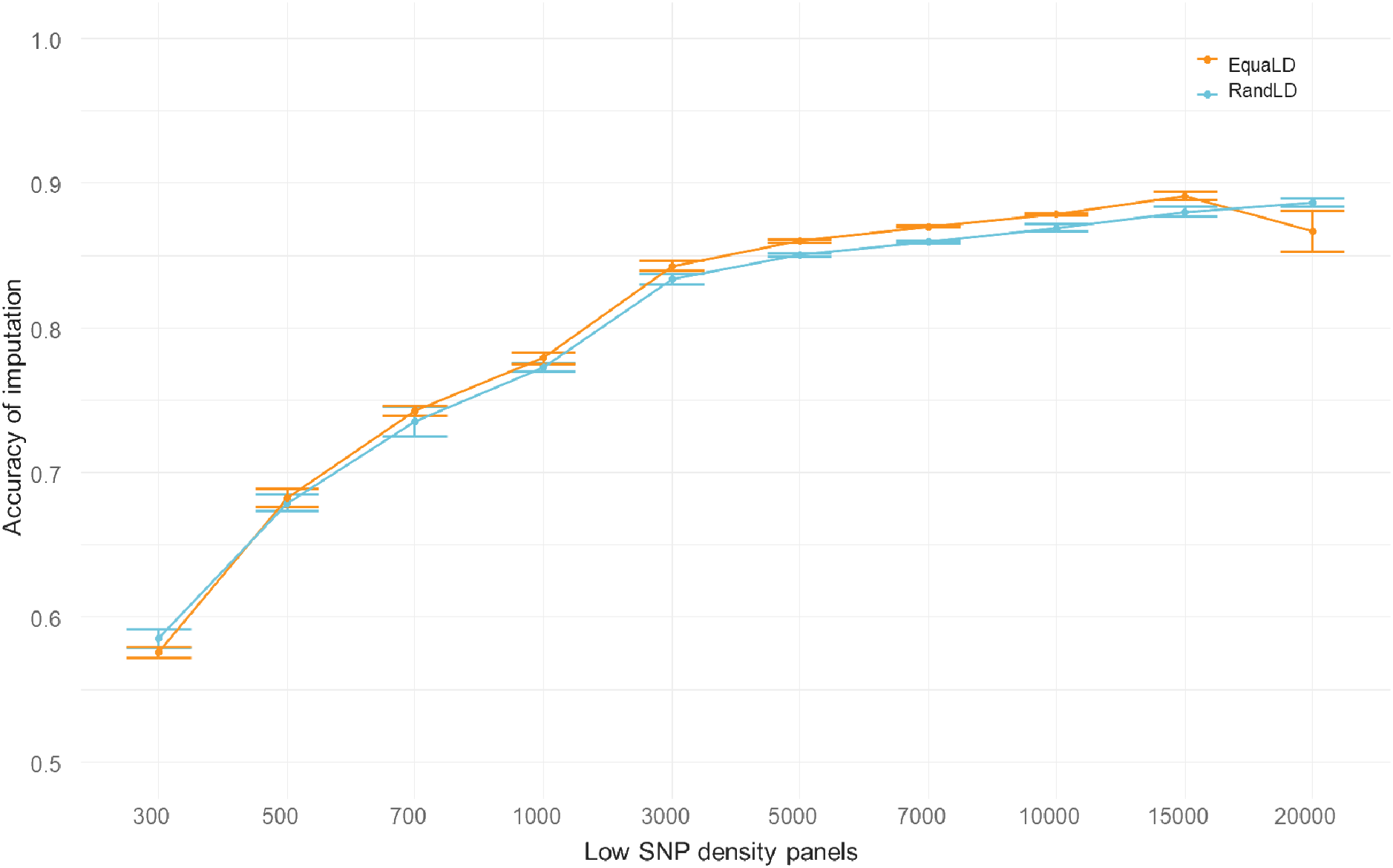
Imputation accuracy of LD-panels imputed to 28K SNPs. Accuracy measured as the Pearson coefficient between true and imputed genotypes for each individual and averaged over the 10 LD-panel replicates for each of the SNP densities. In blue results for the RandLD-panels (randomly sampled), and in orange for the EquaLD panels (equally spaced).

The number of SNPs in each panel post-imputation was slightly lower than in the HD-panel due to the quality controls on the MAF preformed post-imputation. On average, 27K SNPs remained in imputed panels for both SNP selection methods (min=23,941 for 300 SNPs selected using the EquaLD equidistant sampling; max=27,821 for the 20K SNPs RandLD randomly selected).

### Genomic prediction with imputed LD-panels

After imputation, for all starting SNP densities, the accuracy of genomic prediction for both SNP selection methods ranged between 0.63 and 0.65 with a plateau at 0.65 from 3,000 SNPs and above.

At the lowest densities (< 3,000 SNPs, Figure 3 and Table 1), imputation had a positive impact on the accuracy of genomic prediction, with accuracy values similar to those obtained with 3,000 SNPs without imputation. The largest increase in accuracy of genomic prediction due to imputation was observed for the lowest density panel (300 SNPs), where the accuracy of genomic prediction was increased by 11.6% for the RandLD (Table 2) and 7.5% for the EquaLD panels after imputation (Supplementary tables S2 and S3). For SNP densities of 500, 700 and 1,000, imputation increased the accuracy of genomic prediction by 5% on average for the random sampling and 5.5% on average for the equidistant sampling. Both sampling methods had similar performances after imputation.

**Figure 3.**
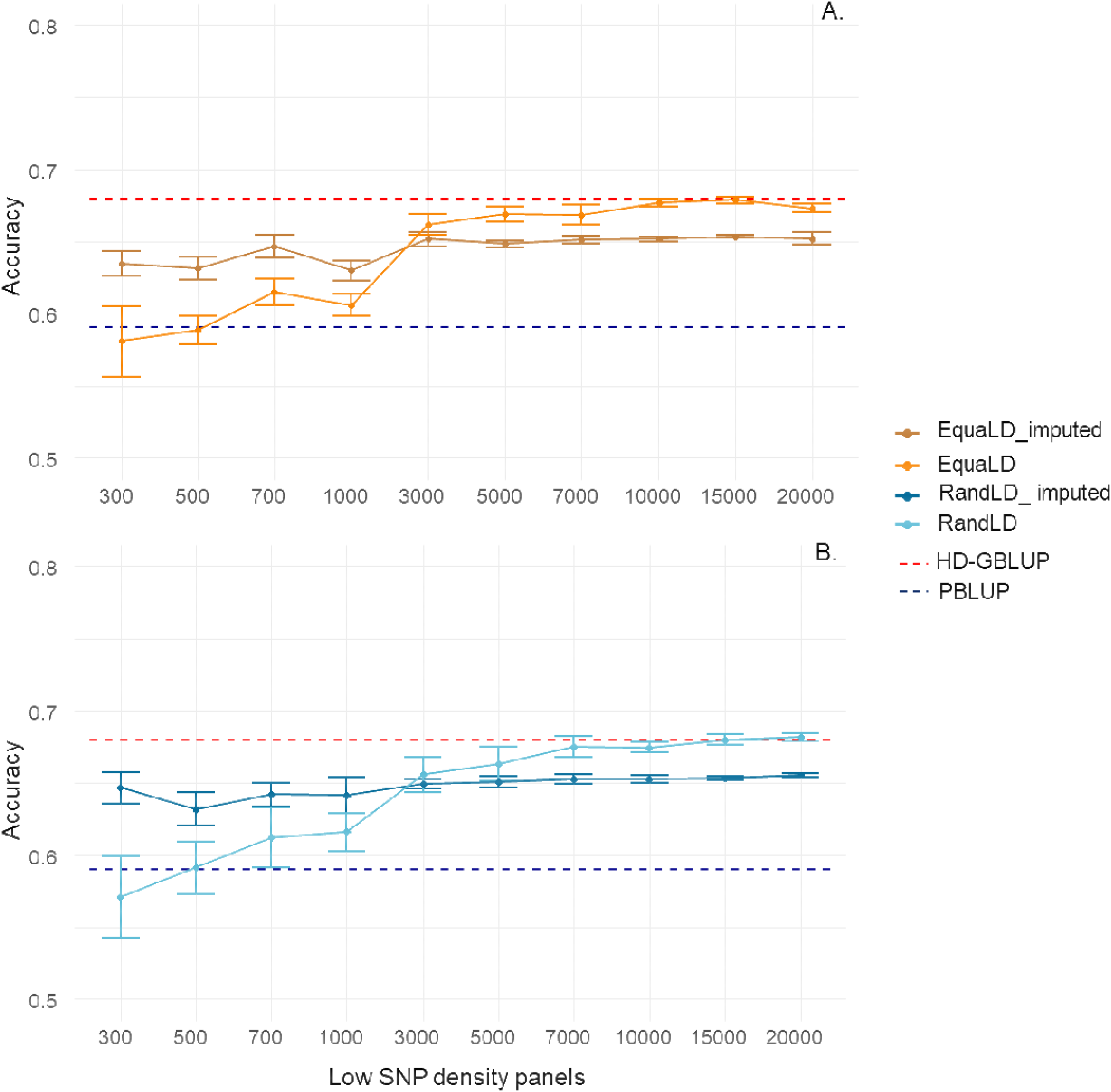
Accuracy of genomic prediction for resistance to *F. columnare* in rainbow trout, obtained with different SNP panels of different densities, before and after imputation, for A. Equally spaced SNPs, and B. Randomly sampled SNPs. The horizontal red dotted line is the average accuracy for the HD-GBLUP (28K) prediction (0.68), the horizontal blue dotted line is the average accuracy for the pedigree-based BLUP prediction (0.59). Figure 3.A. is for equally spaced SNP panels. The orange line is the accuracy value obtained with the equally spaced LD-panels (EquaLD) and the dark orange line is the accuracy obtained after imputation for those panels. Figure 3.B. is for Random SNP panels. The light blue line is the accuracy value obtained with the Random LD-panels (RandLD) and the dark blue line is the accuracy value obtained after imputation for those panels.

**Table 2.** Accuracy of genomic prediction obtained for the random LD-Panels before and after imputation.

| SNP density | Accuracy of RandLD-panel | Accuracy of imputed RandLD-panel | Change in accuracy due to imputation | Difference in accuracy between Imputed LD-panel and HD-GBLUP | Difference in accuracy between Imputed LD-panel and PBLUP | Accuracy of TopLD panels |
| --- | --- | --- | --- | --- | --- | --- |
| **300** | 0.57 ± 0.080 | 0.65 ± 0.078 | +11.6% | -5.0% | 9.5% | 0.62 ± 0.072 |
| **500** | 0.59 ± 0.077 | 0.63 ± 0.078 | +6.3% | -7.1% | 6.9% | 0.66 ± 0.161 |
| **700** | 0.61 ± 0.078 | 0.64 ± 0.077 | +4.7% | -5.6% | 8.7% | 0.63 ± 0.074 |
| **1,000** | 0.62 ± 0.076 | 0.64 ± 0.076 | +4.0% | -5.7% | 8.6% | 0.63 ± 0.074 |
| **3,000** | 0.66 ± 0.074 | 0.65 ± 0.075 | -1.0% | -4.5% | 10.0% | 0.64 ± 0.072 |
| **5,000** | 0.66 ± 0.075 | 0.65 ± 0.076 | -1.9% | -4.3% | 10.2% | 0.65 ± 0.074 |
| **7,000** | 0.68 ± 0.075 | 0.65 ± 0.076 | -3.4% | -4.0% | 10.5% | NA |
| **10,000** | 0.67 ± 0.074 | 0.65 ± 0.075 | -3.4% | -4.0% | 10.5% | NA |
| **15,000** | 0.68 ± 0.075 | 0.65 ± 0.076 | -4.1% | -4.0% | 10.6% | NA |
| **20,000** | 0.68 ± 0.075 | 0.66 ± 0.077 | -4.0% | -3.6% | 11.0% | NA |

RandLD= Random Sampling low-density panels. HD-GBLUP= High density panels (28K SNPs). PBLUP = pedigree-based BLUP. TopLD= Top SNP Low-density panels. Mean of accuracy +/- sd across all 100 values. Values for EqualD in Supplementary tables S2 and S3.

Surprisingly, the accuracy of genomic prediction obtained with 3,000 SNPs was not significantly different before and after imputation, a small decrease of 1% for the random sampling and 1.5% for the equidistant sampling was observed. From 5,000 SNPs or higher densities, the accuracy of genomic prediction obtained after imputation was slightly lower than without imputation (Table 2 and Supplementary table S2 and S3), with on average a decrease of 3% compared to LD-panels. After imputation, there was no or very little bias for all density panels with the average bias ranging between 1.00 and 1.01 (Supplementary Tables S2).

### Cost analysis

To assess the potential to reduce genotyping costs by different genotyping and imputation practices, costs of genotyping and changes in accuracy were estimated for three SNP panels of decreasing densities (57K SNPs, 3K and 300 SNPs) for a breeding population of 8,000 offspring and 200 parents, reflecting the Finnish breeding programmes of rainbow trout. The prices used are estimated for both LD-panels since none of those are available on the market at the moment. For the high-density SNP array (57K SNPs), genotyping is approximately 20€ per sample when genotyping 8,200 fish resulting in a total cost of about 164K €, which can be highly prohibitive for most breeding programmes. Under the assumption that a 3K SNP panel could be genotyped for all 8,200 fish at a cost of 15€ per sample, this would represent a 25% reduction in genotyping cost for a 3% decrease in accuracy compared to the full HD panel. Another possible scenario is that all offspring are genotyped for a very low-density SNP panel (300 SNPs) at a cost of 7.5€ per sample, with parents genotyped for the existing 57K array at a higher cost of 30€ per sample (the price is highly dependent of the number of samples genotyped); in this scenario genotyping would cost 66K€. This reduces the genotyping cost by 60% compared to the price of the 57K SNP for only a 4% decrease in accuracy using the imputation approach. Moreover, *F. columnare* infects small fish, well before they can be individually tagged and identified. Therefore, even for an accurate pedigree-based evaluation, the offspring and parents need to be genotyped in order to recover the pedigree, meaning that the 300 SNP panel could be combined for both parentage assignment and imputation-based genomic selection.

## Discussion

In a previous work (41) we estimated a moderate heritability for resistance to *F. columnare* in this Finnish rainbow trout population (h^2^g=0.21) and we showed that genomic evaluation improved the accuracy of estimated breeding values compared to pedigree-based evaluation. The results obtained in the current study show that the use of low-density panels, along with imputation, results in a higher accuracy of genomic prediction than pedigree-based PBLUP or using LD-panels with no imputation, and could be an efficient way to implement genomic selection.

### Performance of LD-panels

In the current study, no matter the SNP sampling method used to create LD-panels, we showed that 3,000 to 7,000 markers without imputation would result in comparable selection accuracy to the full 28K HD-panel. With only 3,000 SNPs the prediction reached 96.4-97.3% of the HD-panel prediction accuracy, and with 7,000 SNPs between 98.3-99.3% of the HD-panel accuracy was obtained with RandLD and EquaLD panels, respectively. With the panels containing 300 or 500 SNPs, the accuracy was in the range of the one obtained with PBLUP, with a non-significant decrease of accuracy by 3.3% or 1.6% for the 300 panels for RandLD and EquaLD, respectively (Supplementary table S3).

Those values of 3K-7K SNPs being sufficient in aquaculture species to reach near maximum accuracy of prediction are in the range of what has been reported in other aquaculture species for various traits, although the maximum accuracy was usually estimated based on medium density panels ranging from 10K to 70K SNPs (17,18,20,26,29,32,51–54). The number of SNP required to accurately estimate breeding values in aquaculture species and particularly salmonids are substantially lower than what was reported for terrestrial species which ranged between 49K SNPs for pig and 168K SNPs for Holstein cattle (55,56). The high accuracy of genomic prediction obtained for lower density in aquaculture species than terrestrial ones can be explained by the fact that predictions are obtained from close relatives (training population composed of full and half-sib of the validation population) with very high within family linkage disequilibrium as well as long-range linkage disequilibrium.

The high accuracy obtained with such low-density panels in aquaculture populations is most likely because aquaculture breeding programmes rely on large families and sib-testing, thus fish in the training and validation populations have a close relationship. In such populations, many individuals will share long haplotype blocks since they have not been broken by recombination over generations, thus only a few SNPs per chromosome are needed to capture all the genomic information in aquatic species. Furthermore, in salmonids the limited male recombination across most of the genome (57,58) is responsible for a slower decay of linkage disequilibrium and long un-recombined haplotype blocks being shared by individuals. In a previous study on Atlantic salmon, we showed that reducing both SNP density and relationship between training and validation population led to a dramatic decrease in accuracy of genomic prediction (59).

Another possible explanation is the existence of long-range linkage disequilibrium that has been previously characterised in salmonids species. In this rainbow trout population from LUKE, we estimated the linkage disequilibrium between two SNPs about 1Mb apart to be on average 0.11 (± 0.16) (41), lower than the 0.13 and 0.25 values estimated in other rainbow trout populations (51,60). In their study on rainbow trout resistance to BCWD, Vallejo et al., (2018) (51) showed that the accuracy of genomic prediction obtained with only 3K SNPs was almost as good as the accuracy obtained with 45K and partly explained these results by the high level of long-range linkage disequilibrium in this population (r^2^ ≥ 0.25 spanning over 1Mb across the genome). Vallejo et al., (2018) (51) estimated the number of independent chromosome segments, and thus the number of informative SNPs for an efficient GWAS and GS study, to be 20K, however the effective population size (Ne) they estimated was high. Based on the same formula (4LNe) from Daetwyler et al., (2010) (61) and Ne estimated in French rainbow trout populations by D’Ambrosio et al., (2019) (60), the number of independent chromosome segments would vary between 3.1K and 8.8K. The 8.8K are in range of the size of efficient LD-panel we estimated in our study.

Finally, in aquaculture species, the optimum density panel to obtain near maximum accuracy without imputation varies slightly depending on the species and trait. For most aquaculture species a LD-panel of 3,000 SNPs was sufficient to reach an accuracy similar to the HD-panel (9). In Atlantic salmon, depending on the population, between 1K and 5K SNPs were required to reach the same accuracy as the one obtained with 33K or 70K (26,29). Yoshida et al., (2019) (28) showed that with only 3,000 SNPs the accuracy of prediction for rainbow trout resistance to *P. salmonis* was similar to the accuracy obtained with the HD panel of 27K SNPs using a Bayes C approach. Two studies on multiple aquaculture species, (17,18) compared *in* silico LD-panels to HD-panels with a density ranging between 12K to 40K for 5 fish species (common carp, turbot, sea bass, rainbow trout and Atlantic salmon) and reported that LD-panels containing between 3,000 to 10,000 SNPs would be sufficient to obtain near maximum accuracy. Recently similar prediction accuracies with 2,000 SNP and 4,500 SNPs were obtained in flat oyster (20). In European sea bass and sea bream, genotyped initially with about 60K SNPs achieved 90% of the accuracy of the HD-panel with only 6,000 SNPs (19). In our study, as in most previously published studies (17–19,54), further reduction of the density of LD-panels, between 700 and 1,000 SNPs, resulted in a significant drop in the accuracy of genomic prediction but they remained higher than the accuracy obtained with a PBLUP. In the cases of LD-panels with less than 1K SNPs the GEBVs were also more biased (51,54) whereas in the current study the RandLD and EquaLD panels resulted in very little bias. This difference in the performance of LD-panels might also be due to the trait - a lower density panel would perform better for a more heritable trait as simulated by Dufflocq et al., (2019) (30). In rainbow trout, Al-Tobasei et al., (2021) (54) reported that for fillet firmness, with a moderate to high heritability of 0.38, a LD-panel composed of about 1K SNPs would have a similar predictive ability than the HD-panel containing 50K SNPs. However, for fillet yield, with a lower heritability of 0.20, more SNPs (11K) were required to reach a similar prediction accuracy.

Our results confirm that for aquaculture species accurate genomic prediction can be achieved with a low number of markers ranging between 3,000 and 7,000. The long-range linkage disequilibrium and low recombination rate that exist in salmonids and other aquaculture species along with the structure of breeding programmes relying on close relationships between the training and validation population are likely the main drivers for the good performance of low density panels.

### LD-panels based on GWAS results

Previously (Fraslin et al., 2022) (41) we detected a main QTL for resistance to *F. columnare* in this rainbow trout population, which increased the accuracy of genomic prediction when included in a GBLUP approach in which SNPs were weighted by their allele substitution effect. In the current study we wanted to test the effect of including SNPs significantly associated with resistance to *F. columnare* in the LD-panels (TopLD-panels). This strategy significantly increased the accuracy of genomic prediction compared to the RandLD and EquaLD panels for densities of 1,000 SNPs and below. This was expected as the TopLD panels had the SNPs with the highest significant association with resistance, and thus the highest effect. Similarly, Al-Tobasei et al., (2021) (54) reported higher accuracy for genomic prediction of fillet firmness in rainbow trout when using down to 800 SNPs prioritised based on the proportion of genetic variance explained, but with highly inflated predictions. However, in that study the GWAS was performed on the full population including the validation set for the genetic evaluation, which has an important impact on the bias of the predictions. Two studies on rainbow trout resistance to *F. psychrophilum* (51,62) used LD-panels of 70 or 49 SNPs located within previously detected QTL associated to this trait in a previous generation of the same population (63), and showed that these performed as well or even better than HD panels in terms of accuracy, and therefore could be used to accurately predict GEBVs in subsequent generations. Those panels performed better than the LD-panel without the main QTLs (51), highlighting the importance of including SNPs associated with the trait of interest.

In the current study, for densities of 3,000 and 5,000 SNPs, RandLD or EquaLD performed significantly better than TopLD to predict the value of the fish in the validation set. In our population the genetic architecture of resistance was oligogenic, with a main QTL on chromosome Omy3, minor QTL and a polygenic effect (41). As the density increased, SNPs with links to QTL of smaller size were included in the panels, but SNPs from the main QTLs were overrepresented, with a clear oversampling of SNPs from the chromosome Omy5. In a previous work by Calboli et al., (2022) (42) using two rainbow trout populations, we showed that there is a smaller QTL on Omy5 that is spanning 55Mb with a large number of SNPs in very high linkage (r^2^=0.77 on average) (42). This high linkage is responsible for a relatively high effect of all the SNPs in this 55Mb and, as the density of the TopLD panels increases, more SNP from Omy5 with redundant information are sampled, which do not contribute to the accuracy of genomic prediction since no or very little new information is added. The overrepresentation of SNPs from QTLs in the TopLD panels led to highly biased prediction (on average 0.55). Furthermore, creating those LD panels based on GWAS results, also promising for very low densities, would not be applicable in practice. Indeed, not only this analysis requires an extra analysis, the GWAS, to be performed on the training population, but mainly it requires the development of a new LD-panel for each population (and trait) since the QTLs might not be shared between populations (and traits). For resistance to CD, QTLs are shared between two close Finnish populations (42) but not between Finnish and American populations (41,64). The limited use of such LD-panel would potentially increase its cost, and therefore defeat the purpose of using a LD-panel.

### Performance of imputed LD-panels for genomic evaluation

The accuracy of imputation increased rapidly with the number of SNPs in the LD-panel and from 3,000 SNPs upwards, the imputation accuracy was 0.84-0.86 (+/- 0.001-0.003) for RandLD and EquaLD, and it remained under 0.90 even when 20,000 SNPs were included in the LD panel (*i.e*. only about 8,000 missing SNPs to impute). Those values are in the range of what was previously reported for Atlantic salmon by Tsai et al., (2017) (29), and lower than what was achieved by Kijas et al., (2017) (65) or Yoshida et al., (2018) (27) who used larger reference populations for imputation. The higher imputation accuracy obtained in other populations or in cattle could be due to a deeper pedigree in those populations, improving phasing and therefore imputation.

Interestingly, in our study, the accuracy of genomic prediction post imputation for both RandLD and EquaLD panels was quite stable no matter the starting density before imputation. The accuracy of genomic prediction did not seem to be affected by lower imputation accuracies, as observed for the lowest densities (300 – 700 SNPs). In a simulation study on rainbow trout, Dufflocq et al., (2019) (30) also showed that there was no significant differences in the accuracy of genomic prediction obtained after imputation with imputation error rates of 10%, 5% or 1%. In most studies published in aquaculture species the accuracy of genomic prediction after imputation was similar or slightly lower than the accuracy obtained with HD panels. Interestingly, in our study we never reached the accuracy of genomic predictions obtained with the HD-panel, and for densities over 5,000 the accuracy genomic predictions obtained with the LD-panel was significantly higher than the accuracy obtained with the same panel after imputation. A similar observation was made by Vallejo et al., (2021) (62) in their study on rainbow trout resistance to *F. psychrophilum*. They imputed a LD-panel of 7K SNPs to a high-density of 32K SNPs and reported a lower accuracy of genomic prediction after imputation. However, since the actual genotyping was performed with 7K SNPs the accuracy of imputation could not be estimated and this decrease could not be linked to imputation errors.

In order to better understand what could be the cause of this decrease in accuracy post imputation for LD-panels above 5K, we first imputed the HD-panel to get a SNP call rate of 100% (as done in previous studies) (52,66) and re-estimated the accuracy of the imputed-HD-panel. The accuracy of genomic predictions we obtained for the imputed-HD-panel was 0.65 (± 0.077), lower than the HD-GBLUP prior to imputation (0.68 ± 0.076) but not significantly different from the accuracy of the imputed LD-panels. We also performed a second test by setting all the genotypes that were missing in the un-imputed HD-panel to missing in the imputed LD-panel and re-estimating the accuracy of genomic prediction (Supplementary Figure S1). This resulted in a significant increase of the accuracy of genomic prediction compared to the imputed LD-panel, although it remained slightly lower than the accuracy of the un-imputed LD-panel (supplementary Figure S1). These results point towards an important impact of the missing genotypes, which is erased by imputation. In this dataset, 9% of the SNPs that passed the QC controls had a missing rate significantly different between alive and dead fish. Those missing genotypes might provide information that is lost during imputation and thus result in the lower accuracy observed after imputation or they might generate bias and inflate the accuracy that is corrected by the imputation.

### Cost efficiency of genotyping strategies

In this study we showed that using a LD-panel to genotype rainbow trout and perform a genomic prediction of resistance to CD would result in a minor reduction of the accuracy of prediction (3%) compared to the use of a HD-panel for a considerable reduction in cost (about 25%). However, the price of 15€ per sample for genotyping using a 3K SNP panel is hypothetical as no such panel exists for rainbow trout, and in reality this LD-panel could be more expensive than estimated, thus reducing the interest of LD-genotyping. One solution would be to incorporate those 3K SNPs on a multispecies SNP panel that would be produced in more volume and thus be less expensive. Such multi-species panels have been developed for various aquaculture species (*Sparus aurata* and *Dicentrarchus labrax* ((67), *Crassostrea gigas and Ostrea edulis* (68), or *Colossoma macropomum* and *Piaractus mesopotamicus* (69)). An interesting solution would be to develop a very low-density SNP panel or use targeted-GBS to genotype 300 to 500 SNPs (to account for a decrease of the number of SNPs post QC) and combine it with imputation using HD-genotyped relatives as reference population. Such very LD-panel could be developed to be specific to a population, could include SNPs located within QTLs associated to traits of interest and be used not only for genomic selection but also for parentage assignment. In breeding programmes, the number of traits in the selection index is quite high and including QTLs associated with all of them would not be possible in very-low-density panels. However, most of the traits are polygenic and there are very few traits of interest that are controlled by a major QTL (70). The careful design of very LD-panels (equally spaced SNPs along all chromosomes) will maximise imputation accuracy and create affordable LD-panels that are highly efficient for genomic selection when combined with imputation.

## Conclusion

In conclusion, the use of low-density SNP panels may reduce the costs of genomic selection in rainbow trout without a major reduction in the prediction accuracy of breeding values. Using low-density SNP panels (about 3,000 markers) or very low-density SNP panels (about 300 markers) combined with imputation using HD-genotyped parents would result in a decrease of only 3-4% compared to a HD-genotyped population, which corresponds to an increase of 10.5% to 11% compared to a pedigree-based prediction. The good performance of such low-density panels might be potentially valid for most aquaculture species with long-range linkage disequilibrium, low recombination rate and breeding programmes reliant on sib-testing. Our findings suggest that a cost-effective genomic evaluation to improve resistance to *F. columnare* in rainbow trout is feasible and LD genotyping along with imputation could be a way to speed-up the implementation of genomic selection in low or medium scale breeding programmes.

## Acknowledgment

Lotta Mäkinen, Heikki Koskinen, Antti Nousiainen, Miika Raitakivi, and the skilled staff of Savon Taimen Oy and Hanka Taimen Oy are thanked for their expertise in data collection and fish rearing.

## Author contribution

Clémence Fraslin: Formal analysis, Data curation, Visualization, Writing – original draft, Writing – review & editing. Diego Robledo: Supervision, Project Administration, Writing – review & editing. Antti Kause: Conceptualization, Resources, Supervision, Project administration, Funding acquisition, Writing - review & editing. Ross D. Houston: Conceptualization, Resources, Supervision, Project administration, Funding acquisition, Writing – original draft, Writing – review & editing.

## Supplementary material

**Supplementary Table S1. Number of SNPs in low-density panels**

RandLD= Random Sampling of SNPs in the low-density panels. EquiLD = Equidistant sampling of SNPs in the low-density panels.

**Supplementary Table S2. Values of accuracy and bias of genomic prediction obtained for the random, equidistant and top LD-Panels before and after imputation.**

RandLD= Random Sampling of SNPs in the low-density panels. EquiLD = Equidistant sampling of SNPs in the low-density panels. TOP-LD= Top SNP Low-density panels. Mean of accuracy +/- sd across all 100 values.

**Supplementary Table S3. Proportion of increase or decrease of the accuracy of genomic prediction obtained for the random, equidistant and top LD-Panels before and after imputation compared to pedigree and HD-panels.**

RandLD= Random Sampling of SNPs in the low-density panels. EquiLD = Equidistant sampling of SNPs in the low-density panels. TOP-LD= Top SNP Low-density panels. Mean of accuracy +/- sd across all 100 values. PBLUP= pedigree-based BLUP. HD-GBLUP= genomic based BLUP obtained with the High density (HD) panel.

**Supplementary Figure S1.**
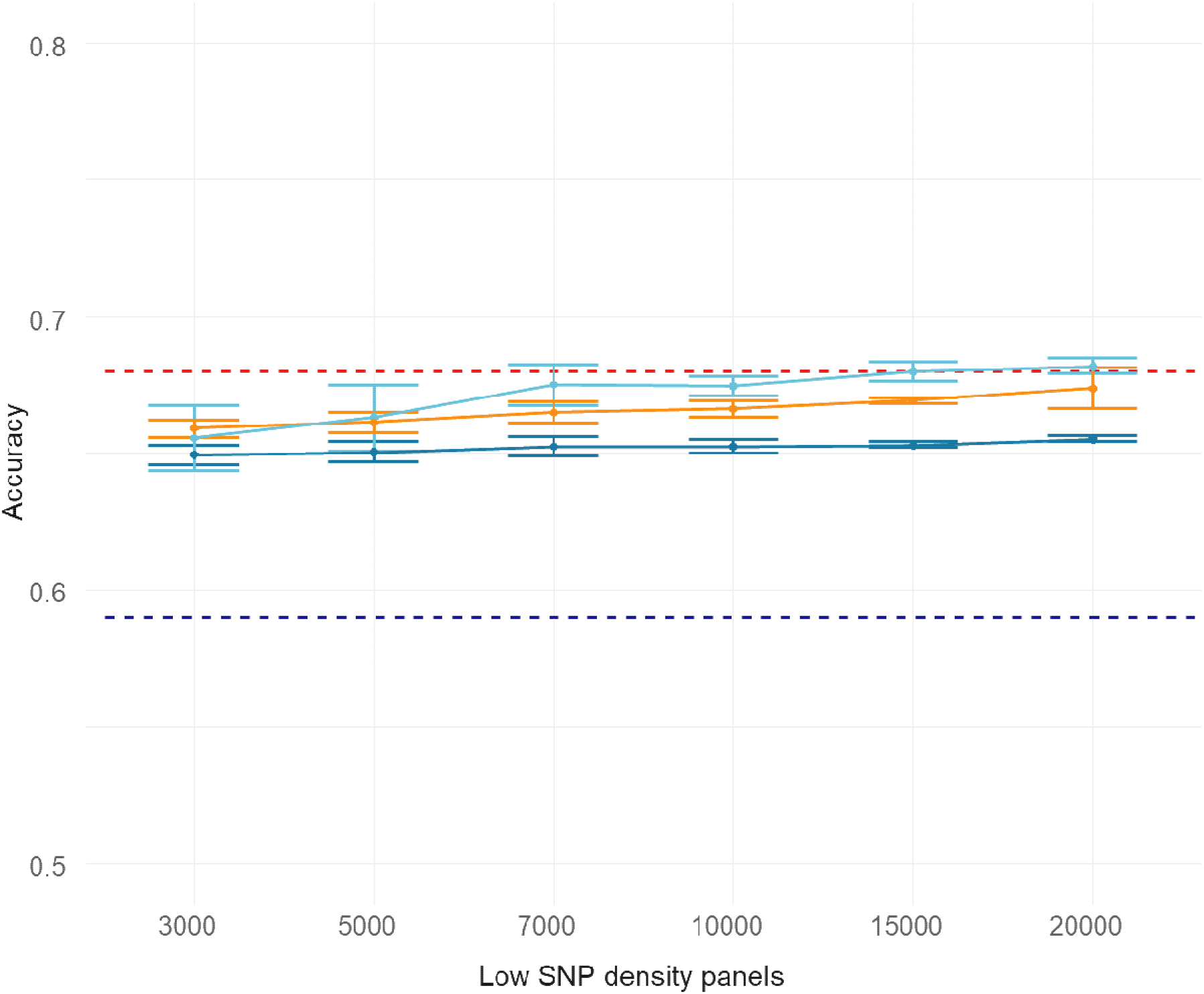
Accuracy of genomic prediction for resistance to *F. columnare* in rainbow trout, obtained with SNP panels of different densities, before and after imputation and before or after re-setting genotype missing in the HD-panel as missing after imputation. The red dotted line is the average accuracy for the HD-GBLUP (28K) prediction (0.68), the blue dotted line is the average accuracy for the pedigree-based BLUP prediction (0.59). The LD panels were created with random SNP sampling (RandLD). The blue line is the accuracy value obtained with the LD-panels (RandLD) and the dark blue line is the accuracy value obtained after imputation for those panels. The orange line is the accuracy obtained after imputation of those panels and after re-setting all the missing genotype from the HD-panel as missing in the imputed-LD-panels.

